# *DPYSL2*/*CRMP2* isoform B knockout in human iPSC-derived glutamatergic neurons confirms its role in mTOR signaling and neurodevelopmental disorders

**DOI:** 10.1101/2022.11.19.517191

**Authors:** Kyra L. Feuer, Xi Peng, Christian Yovo, Dimitri Avramopoulos

## Abstract

*DPYSL2/CRMP2* is a microtubule-stabilizing protein crucial for neurogenesis and associated with numerous psychiatric and neurodegenerative disorders. *DPYSL2* has multiple RNA and protein isoforms, but few studies have differentiated between them or explored their individual functions. We previously demonstrated in HEK293 cells that a schizophrenia -associated variant in the *DPYSL2* B isoform (*DPYSL2*-*B*) reduced the length of cellular projections, created a transcriptomic disturbance that captured schizophrenia etiology, and was acted upon by the mTOR pathway. In the present study, we follow up on these results by creating, to our knowledge, the first models of endogenous *DPYSL2*-*B* knockout in human induced pluripotent stem cells and excitatory glutamatergic neurons. We use CRISPR/Cas9 to specifically knock out *DPYSL2*-*B* and observe corresponding reduction of its RNA and protein. The average length of dendrites in knockout neurons was reduced up to 58% compared to controls. Transcriptome analysis reveals disruptions in pathways highly relevant to psychiatric disease including mTOR signaling, cytoskeletal dynamics, immune function, calcium signaling, and cholesterol biosynthesis. We also observed a significant enrichment of our differentially expressed genes in schizophrenia GWAS-associated loci. Our findings clarify the functions of the human *DPYSL2*-*B* isoform and confirm its involvement in molecular pathologies shared between many psychiatric diseases.

## INTRODUCTION

Dihydropyramidase-like 2 (*DPYSL2*; also known as *CRMP2/TOAD-64/Ulip2/DRP-2/unc33/CRMP62/TUC*-2) is a ∼140kb gene on chromosome 8p21.2 that encodes collapsin response mediator protein 2 (*CRMP2*). *DPYSL2* belongs to a family of homologous genes (*DPYSL1*-*5*/*CRMP1*-*5*) that are conserved across vertebrates and are highly expressed during early brain development in neurons and glia ^1; 2^. *DPYSL2*/CRMP2 is the most well-studied of the CRMPs and was first identified as an intracellular respondent of Semaphorin 3A, a repulsive axon guidance molecule in neurogenesis ^3^. CRMP2 has since been realized to have diverse functions within the cytosolic ecosystem. It is most well-known as a microtubule-stabilizing protein that functions by forming tetramers with itself or other CRMPs that stabilize assembled microtubules and traffic tubulin dimers to their plus end ^4^. Consequently, CRMP2 is a mitigator of diverse cytoskeletal-related functions including mitosis, cell migration ^5; 6^, endocytosis, mitochondrial morphology and motility ^7^, kinesin and dynein-facilitated molecular transport, cell polarity, and dendritic and axonal elongation ^2; 8; 9^. Additionally, CRMP2 acts as a modulator of calcium homeostasis and neurotransmitter release by both regulating the trafficking of calcium channel subunits and binding to NMDAR receptors to inhibit their activity ^2; 10; 11^. Recent studies have even found that post-transcriptional modifications of certain CRMP2 isoforms uncover nuclear localization sequences that allow their proteins to enter neuronal nuclei and potentially function as transcription factors ^12; 13^. CRMP2’s functions are dynamically regulated by post-translational modification of its C-terminus. In particular, phosphorylation at various amino acids by the CDK5/GSK3β and Rho pathways affects CRMP2’s affinity for assembled microtubules and tubulin dimers, which subsequently results in fluid elongation or retraction of axons ^2^. CRMP2 activity is also regulated by the mTOR kinase, which mediates translation efficiency of CRMP2 mRNA by phosphorylating translation initiation proteins that bind to a 5’ Terminal OligoPyrimidine (TOP) motif in the 5’ untranslated region (UTR) ^14-16^.

Given CRMP2’s role in neurogenesis, it’s no surprise that its dysfunction has been documented in numerous psychiatric and neurological disorders ^2; 17^. Genetic variants in *DPYSL2* have been associated with schizophrenia (SCZ) ^14; 18; 19^, bipolar disorder (BPD) ^18^, Alzheimer’s disease (AD) ^20^, autism spectrum disorder (ASD) ^21; 22^, and intellectual disability ^23^. Mouse *DPYSL2* knockout models recapitulate SCZ and ASD-associated pathologies including behavioral abnormalities, gross structural changes in brain, neuron morphological abnormalities, deficits in synaptic pruning, and perturbation of glutamatergic signaling ^2; 5; 8; 24; 25^. Reduced CRMP2 abundance has been observed in postmortem brain samples of patients with SCZ, BPD, major depressive disorder (MDD) ^26^, and Down’s syndrome ^27^. Hyperphosphorylation of specific residues of CRMP2 has been observed in mouse models of multiple sclerosis and amyotrophic lateral sclerosis, and suppression of this modification was found to ameliorate pathologies ^28; 29^. In AD, CRMP2 shares much of the same fate as fellow microtubule-associating protein Tau: it is regulated by the same kinases (Cdk5/GSK3β)^30-32^, becomes hyperphosphorylated ^33^, and eventually associates with neurofibrillary Tau tangles ^34^. The ratio of phosphorylated: unphosphorylated CRMP2 in blood has been found to be decreased in SCZ and increased in lithium-responsive BPD ^35^.

Despite the large amount of interest and research on CRMP2, few studies have differentiated between its isoforms. The human *DPYSL2* gene has 16 exons which are alternatively spliced into multiple transcripts, three of which have a consensus coding sequence: *DPYSL2*-*A*/*CRMP2-A* (ENST00000521913/ NM_001197293); *DPYSL2*-*B*/*CRMP2*-*B* (ENST00000311151/NM_001386), and *DPYSL2*-*C*/*CRMP2*-*C* (ENST00000523027/NM_001244604). These three isoforms share most of their sequence but are differentiated by alternative first exons, and most of what we know is restricted to *DPYSL2*-*A* and *DPYSL2*-*B. DPYSL2*-*B* is considered the canonical transcript because it was discovered first and is the most conserved across species ^2; 3; 36; 37^. Many studies of CRMP2 reference the *DPYSL2*-*B*/CRMP2 gene sequence/mRNA/cDNA/protein; but they either don’t consider the existence of alternate isoforms or their experimental manipulations involve regions of *DPYSL2*/CRMP2 that are common to multiple isoforms ^2; 38; 39^.

However, a few studies in chick and mouse have shown that while CRMP2-B can localize to the soma, dendrites, and less frequently axons, CRMP2-A is usually present in just the axons and soma ^36; 37; 40^. One of these studies found that overexpression of CRMP2-B resulted in increased branching but decreased length of axons, while overexpression of CRMP2-A did the opposite; suggesting CRMP2-A and CRMP2-B have opposing effects and work in tandem to regulate axonogenesis ^36^. Additionally, recent studies have implicated CRMP2-A as a molecular brake on carcinogenesis. Downregulation of CRMP2-A has been observed in prostate cancer biopsies, and deletion of CRMP2-A in human lung adenocarcinoma cells results in cytoskeletal reorganization and progression of the epithelial-mesenchymal transition ^41^, a process in which epithelial cells acquire mesenchymal properties that enhance metastasis ^42^. CRMP2-A has also been shown to regulate the differentiation of hematopoietic stem cells to monocytes, and stimulation of the KLF4-CRMP2-A pathway as a candidate therapeutic for acute myeloid leukemia has been suggested ^43; 44^.

In previous work we reported a SCZ-associated polymorphic CT dinucleotide repeat (DNR) at the 5’-UTR of *DPYSL2-B*. This allele was found to respond to mammalian target of Rapamycin (mTOR) signaling, and to show allelic differences in reporter assays ^19^. Following up on that work we later introduced this repeat to HEK293 cells and observed a significant reduction of the corresponding CRMP2-B isoform, shortening of the HEK293 cellular projections, and marked differences in transcriptome analysis. The changing transcriptome also showed a reverse expression signature from that seen with antipsychotic drugs from ConnectivityMap (CMAP) and other SCZ gene perturbations supporting the importance *DPYSL2*-*B* and a role for mTOR signaling in SCZ ^14^. We now seek further support for our conclusions in a better model system.

In the present study, we used CRISPR/Cas9 in human induced pluripotent stem cells (iPSCs) to create, to our knowledge, the first knockout of the endogenous *DPYSL2-B* isoform in human cells. We then differentiated these edited iPSCs into excitatory glutamatergic neurons and investigated the impact of *DPYSL2*-*B*/CRMP2-B knockout at the transcript, protein, and morphological levels.

## METHODS

### sgRNA Design and Cloning

Two sgRNAs were designed to create a 505bp deletion targeting the first exon (ENSE00002094501) of the *DPYSL2* B isoform (ENST00000311151). sgRNA oligos with BbsI sticky ends (IDT, 25nmole standard desalted, **Supplementary Table 1**) were ligated into the pDG459 plasmid (Addgene 100901) as previously described ^45^. 5uL of ligation reaction was transformed into 25uL of One Shot TOP10 Chemically Competent E.Coli (ThermoFisher C404003) following the manufacturer’s recommendation. Transformed bacteria were plated on LB agar plates with 100ug/ml carbenicillin and incubated at 37C overnight. The following day colonies were picked and cultured in 5mL of Luria Broth with 100ug/mL carbenicillin overnight. Plasmids were isolated from 3mL of the cultures (Qiagen miniprep kit 27104) and screened for correct insertion via Sanger sequencing with one primer for each insertion site (IDT, 25nmole standard desalted, **Supplementary Table 1**).

**Table 1.**
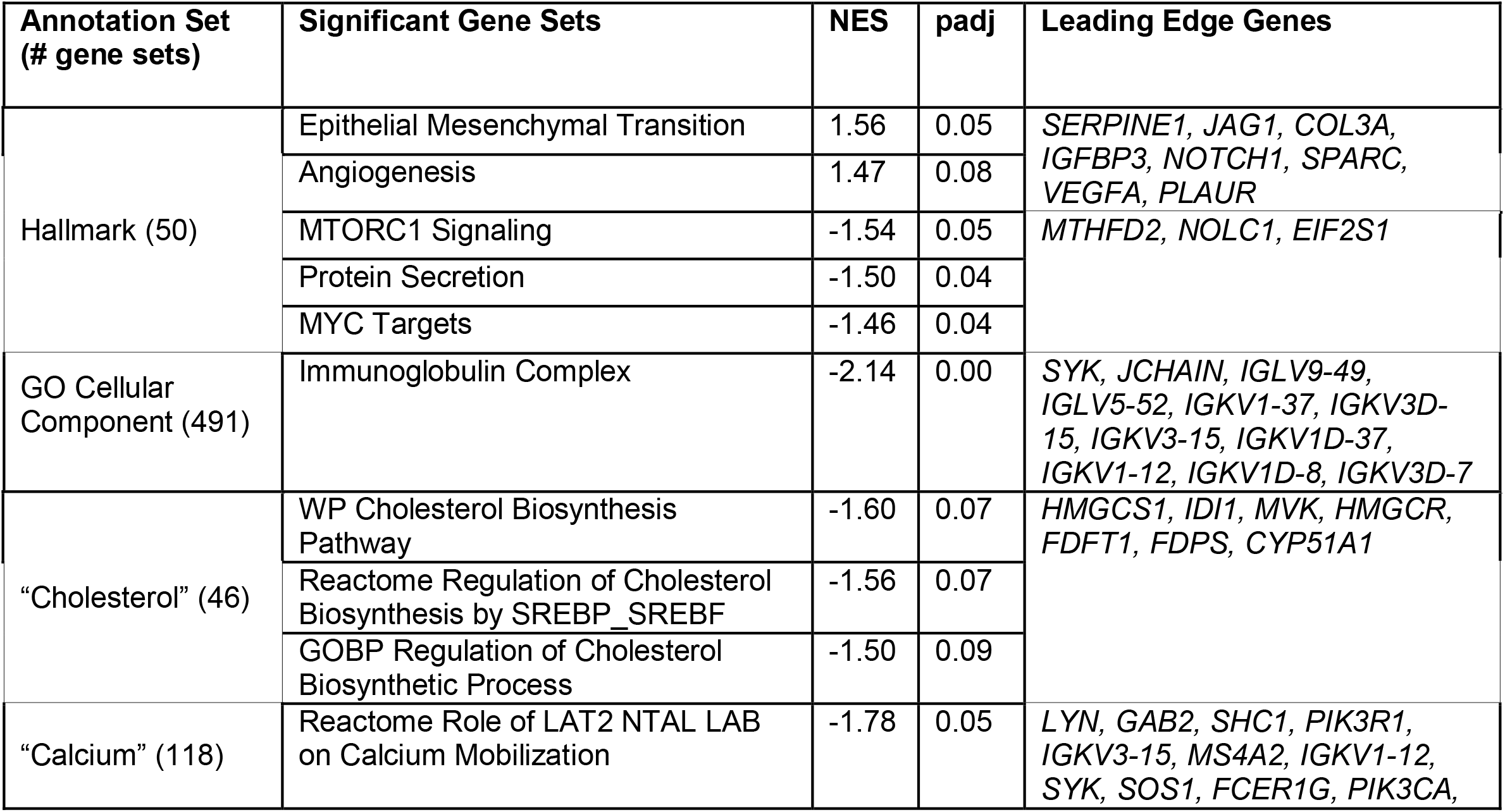
Gene Set Enrichment Analysis Results. Annotation sets represent groups of gene sets related to a biological category. Those within quotation marks are custom, hypothesis-driven sets created by searching MSigDB for gene sets with the quoted term. Significant gene sets are those that were differentially expressed with an padj < 0.1. The Normalized Enrichment Score (NES) indicates the degree to which that gene set was enriched for differential expression in the frameshift neurons, with a negative score indicating downregulation and a positive score indicating upregulation. The Leading Edge Genes are the genes with the largest contribution to the NES. For the annotation sets with one significant gene set (GO cellular component and calcium), the listed leading edge genes are those that contributed to the NES for that individual gene set (aka the core enrichment set). For the Hallmarks and Cholesterol sets where multiple gene sets were significant, the listed leading edge genes are those in common between all the gene sets (aka were shared between the core enrichment sets for all significant gene sets within that annotation set).

### iPSC Quality Testing, Culture and Maintenance

The iPSC line (MH0185922, African male control) was obtained from the NIMH/RUCDR stem cell repository (https://www.nimhgenetics.org/). Upon receipt the cell line was authenticated and screened for chromosomal abnormalities with the Illumina Infinium Global Screening Array-24 v2.0 by the Johns Hopkins Genetic Resources Core Facility. Mycoplasma screens were negative. For culture, all plates were coated with 5ug/mL laminin (Biolamina LN521) in DPBS with magnesium and calcium (Quality Biological 114-059-101) and incubated at 37C for at least 2 hours. iPSCs were cultured feeder-free on the prepared plates in StemFlex media with supplement (Gibco A3349401) at 37°C with 5% CO_2_ unless otherwise specified. When cells reached 70-80% confluency they were dissociated with Accutase (MilliporeSigma A6964) and passaged ∼1:6; or frozen in StemFlex with 10% DMSO at – 80°C for 1-7 days before being transferred to liquid nitrogen for long-term storage. On days of thawing, passaging, or transfection the media was supplemented with 10uM Rock Inhibitor (Y-27632 dihydrochloride, Tocris 1254).

### CRISPR/Cas9 transfection, screening, and single-cell cloning of iPSCs

24 hours before transfection, iPSCs were plated on 24-well plates at 5×10^5^ cells per well. 24 hours later, the media was changed to Opti-MEM (ThermoFisher 31985070) with 10uM Rock inhibitor, and the cells were transfected with 500ng of pDG459 and 2uL of Lipofectamine Stem (ThermoFisher STEM00003) per well following the manufacturer’s instructions. One negative control well received Lipofectamine but no DNA. 24 hours after the start of transfection the media was changed to StemFlex with 10uM Rock inhibitor for recovery. 16 hours later the media was changed to StemFlex with 1ug/uL puromycin to select for transfected cells. 24 hours later all cells in the negative control well were dead, indicating that selection was complete. The media was changed to StemFlex without puromycin and the cells were fed every day until confluent and shedding into the media.

To genotype the “bulk” transfection cells, the media from each well was separately collected and spun down. The supernatant was removed, 20uL of Quick Extract buffer (Lucigen QE0905T) was added, and DNA extraction proceeded according to the manufacturer’s recommendations. 5uL of DNA was used in a PCR with the Accuprime PCR kit (ThermoFisher 12339016) supplemented with 10x PCRx Enhancer (ThermoFisher 11495017). Primer sequences are in **Supplementary Table 1**. PCR products were electrophoresed at 120V for 1 hour in 2% agarose gels and imaged with a BioRad GelDoc EZ Imager with Image Lab Software.

For single-cell cloning, bulk transfection wells positive for the deletion were sparse plated in 6-well plates at a density of 500 cells per well and grown for 7-10 days to form distinct colonies. Half of each colony was transferred to a 96-well plate for continued culture. The other half was transferred to a PCR tube with 10uL of Quick Extract buffer and screened as described above. Colonies appearing positive for the deletion by PCR and gel were sent for Sanger sequencing for confirmation with the primers used for PCR.

Re-genotyping of positive deletion clones after initial expansion revealed emergence of a band at the expected size for unedited DNA. To purify the deletion clones, we sparse plated and picked the clones multiple consecutive times. Each time, the “unedited” band reappeared after passaging. Sequencing revealed that this DNA was not unedited, but contained a 1bp insertion in the first exon at the Cas9 cut site of the sgRNA3 protospacer. After confirming that this insertion should cause a frameshift, we used the same single-cell cloning and genotyping protocols to isolate 6 clones with the insertion for further analyses.

### Off-target analysis and Global Diversity Array of frameshift and control iPSCs

Off-target screening was prioritized to candidate sites that were most likely to be edited and/or in functional sequence. A list of 45 candidate off-target sites for the two sgRNAs was obtained from the CRISPOR sgRNA design tool (http://crispor.tefor.net/) and filtered down to those with 3 or fewer mismatches with the on-target protospacer. The resulting sites were run through a Perl script that identified those in exons or overlapping with open chromatin peaks (DNase I hypersensitivity peaks from the encode database; www.encodeproject.org, file name hg38_wgEncodeRegDnaseClustered.txt). This analysis revealed only a single off-target candidate site at chr14:67889176-67889198. The six frameshift clones and the untransfected iPSC line were genotyped for this site with primers listed in **Supplementary Table 1**.

After *Ngn2*-rtTA transduction (see below), the 6 frameshift clones and 6 untransfected control clones were again authenticated and screened for chromosomal abnormalities via the Illumnia Infinium Global Diversity Array by the Johns Hopkins Genetic Resources Core Facility. No chromosomal abnormalities were found. Identity by state analysis of the queried SNPs with plink ^46^ found the 12 lines were 99.9416-99.9537% identical, within the array error rate.

### *Ngn2*-rtTA transduction of frameshift and control iPSCs

iPSCs were transduced with *Ngn2* and rtTA lentiviruses from either Cellomics (CLVP-109-200UL) or the Children’s Hospital of Philadelphia Clinical Vector Core (LV260/LV259) (Supplementary Table 2). The untransfected iPSC line was transduced six individual times to create six control “clones”. Each clone was plated in one well of a laminin-coated 6-well plate in StemFlex with 10uM Rock inhibitor at a density of 2.5 × 10^5^ cells per well. Once the cells were ∼50% confluent (after 2-3 days) the media was changed to StemFlex with 1ug/uL polybrene (Santa Cruz SC-134220), and 10uL each of *Ngn2* and rtTA virus was added to each well. After a 6-hour incubation at 37C the media was changed to fresh StemFlex with 10uM Rock inhibitor. Once the cells were 80-90% confluent (24-48 hours later) the cells were passaged 1:6 and grown until confluent. Five of the six wells were frozen down and the remaining well was passaged for differentiation (see below).

### *Ngn2*-induced differentiation of frameshift and control iPSCs into glutamatergic neurons

To make glutamatergic neurons for RNA-seq and Western Blot, six frameshift clones and six control clones were differentiated across 7 batches (**Supplementary Table 2**). On Day In Vitro (DIV) -2, each iPSC clone was plated across four laminin-coated 6-well plates in StemFlex with 10uM Rock inhibitor at a density of 2.5 × 10^5^ cells per well. On DIV –1, cells were fed with StemFlex. On DIV 0, differentiation was induced by feeding with iN-N2 media {DMEM/F12 with L-Glutamine and HEPES (Thermofisher 11330-032) supplemented with 1.5mg/mL D-Glucose (MilliporeSigma G8270), 0.1% Beta-mercaptoethanol (Thermofisher 21985023), 100ug/mL Primocin (Invivogen ant-pm), 1% N2 (Thermofisher 17502048), 10ng/mL BDNF (Peprotech AF-450-02), 10ng/uL NT3 (Peprotech 450-03), 200ng/mL Laminin (MilliporeSigma L2020), and 2ug/mL Doxycycline (MilliporeSigma D9891)}. On DIV 1, cells were fed with iN-N2 media with 1ug/mL puromycin (MilliporeSigma P8833) to kill any untransduced cells. On DIV 2, 6-well plates were coated with a solution of 1:35 Matrigel (Corning 354230) in DMEM/F12 and incubated at 37C for at least 1 hour. The surviving cells were dissociated with Accutase, neutralized with 1xPBS, and counted with a Countess cell counter. 1×10^6^ cells were plated per well in the prepared Matrigel plates in iN-NBM/B27 media {Neurobasal Medium without L-glutamine (Thermofisher 2110304) supplemented with 2% B27 (Thermofisher 17504044), 1% Glutamax (Thermofisher 35050061), 1% PenStrep (Thermofisher 15140122), 6ug/mL D-Glucose, 10ng/uL BDNF, 10ng/uL NT3, 200ng/mL Laminin, and 2ug/mL Doxycycline}. On DIV 4, 50% of the media was exchanged with iN-NBM(A)/B27 + Dox media {Neurobasal A medium without glucose or sodium pyruvate (Thermofisher A24775) supplemented with 2% B27, 1% Glutamax, 1% PenStrep, 1.1ug/mL D-Glucose, 0.9ug/mL Sodium Pyruvate (Millipore Sigma P5280), 10ng/uL BDNF, 10ng/uL NT3, 200ng/mL Laminin, and 2ug/mL Doxycycline} with 4uM AraC (MilliporeSigma 1162002-250MG) to kill any non-neuronal cells. Starting on DIV 6, the media was exchanged 50% once every 2 days with iN-NBM(A)/B27 + Dox. Starting on Day 12, the media was exchanged 50% once every three days with the same media but without doxycycline. On DIV 21, the neurons were harvested onto ice for immediate RNA extraction or frozen at –80C for Western Blot.

To make glutamatergic neurons for dendrite morphology analysis, three frameshift clones and three control clones were differentiated on a layer of primary rat astrocytes to encourage the cells to spread out. One week before differentiation, primary rat astrocytes were thawed into T75 flasks and fed once every 2-3 days with glial media {(MEM (Thermofisher 11095080) supplemented with 5% FBS (Gemini 900-108), 1% Penstrep, 20mM D-Glucose, 2.5mM L-Glutamine (Thermofisher 25030081), and 0.2mg/mL sodium bicarbonate (Gibco 25080-094)}. On DIV –1 of neuronal differentiation, four-chamber slides (ThermoFisher 62407-294) were coated with 300uL per chamber of a 1:4 dilution of Poly-L-Ornithine Solution (0.01%, Sigma-Aldrich P4957) in Ultrapure water and incubated at 37C for 4 hours. The slides were rinsed twice with Ultrapure water, then coated with a 1:50 solution of laminin in HBSS (Gibco 14025092) and incubated at 37C overnight. On DIV 0, 9 x10^4 rat astrocytes were plated per chamber in the slides in glial media. On DIV 2, the glial media was removed and 1.5 x10^5 neurons were plated per chamber on top of the astrocytes in iN-NBM/B27 media. Differentiation then proceeded as described above except the iN-NBM/27 media was replaced with Neuronal Media {Neurobasal Medium supplemented with 2% B27, 1% Glutamax, 1% PenStrep, 1mg/mL ascorbic acid, 1uM cAMP, 10mg/mL BDNF, 10mg/mL GDNF, 2ug/mL Doxycycline, and 200ng/mL laminin}. Doxycycline was not added starting on Day 12. Immunocytochemistry was performed on DIV21 (see below).

### RNA extraction of iPSCs and glutamatergic neurons

RNA was extracted from pellets of ∼2 million iPSCs or ∼6 million neurons immediately after harvesting with the Quick-RNA Miniprep Kit (Zymo R1054) following manufacturer’s instructions except for two alterations. After adding lysis buffer, the pellets were vortexed for 30 seconds to mechanically lyse cells; and RNA was eluted in 30uL of water instead of 50uL to increase concentration.

### qRT-PCR of *DPYSL2-B* in iPSCs

cDNA was generated from 2 control iPSC clones and 4 frameshift iPSC clones with the Superscript III kit (Thermofisher 18080-051) following manufacturer’s recommendations using 1ug of RNA per sample and random hexamer primers. cDNA was diluted 1:10 in UltraPure water (Invitrogen 10977-015) and used for qRT-PCR with iTaq Universal SYBR Green Supermix (Biorad 172-5121) with primers targeting the first exon of *DPYSL2*-*B*. Primers targeting GAPDH and B-actin were used for loading control reactions. Samples run on a CFX Connect with CFX Maestro software.

### Western Blot of iPSCs and glutamatergic neurons

iPSC lysates were prepared by thawing frozen cell pellets (∼2 million cells each) on ice for 5-10 minutes, then adding cold RIPA Buffer supplemented with 1mM PMSF (Cell Signaling Technologies 8553S) at a ratio of 160uL per 1 million cells. Samples were rotated at 4°C for one hour, followed by centrifugation at 21,000xg for 30 minutes at 4°C. The cleared lysates were transferred to prechilled tubes, quantified with a BCA analysis kit (Thermofisher 23225) and stored at –80C until ready for use. 20ug of protein was combined with 11.25uL of XT Sample Buffer (Bio-Rad 1610791), 2.25uL XT Reducing Agent (Bio-Rad 1610792), and water to a final volume of 45uL. Samples were boiled at 95C for 5 minutes, then cooled on ice for 10 minutes before electrophoresis.

Neuron lysates were prepared by adding cold Phosphosafe Extraction Reagent (MilliporeSigma 71296-3) supplemented with PhosStop (MilliporeSigma 4906845001) and cOmplete protease inhibitor (Sigma 11697498001) to frozen neuron pellets (∼6 million cells each) on ice at a ratio of 100uL per 1 million cells. Once thawed, pellets were incubated on ice for 15 minutes with vortexing every 5 minutes, followed by centrifugation at 21,000xg for 15 minutes at 4°C. The cleared lysates were transferred to pre-chilled PCR tubes, then immediately neutralized by adding 1/3 volume XT Sample Buffer and heating in a thermal cycler to 98C for 10 minutes, then 20C for 1 minute. Neutralized lysates were stored at -20C until ready for use. Neutralized lysates were thawed and combined with XT Reducing Agent, then heated in a thermal cycler at 65C for 10 minutes and 20C for 1 minute before electrophoresis.

Samples were loaded into XT Tricine gels (Biorad 3450129), along with 6uL of EZ-Run Pre-Stained *Rec* Protein Ladder (Fisher BP3603-500) and 3uL of Blue Prestained Protein Standard (NEB P7718S). Gels were electrophoresed in XT Tricine Running Buffer (Bio-Rad 1610790) at 93V for 30 minutes, then 80V for 75 minutes, then 120V for 75 minutes. Protein was transferred onto low-fluorescence PVDF membrane (BioRad 162-0262) via semi-dry transfer with the BioRad Trans-Blot Turbo system on the mixed MW (Turbo) program. After transfer, the membrane was quickly washed with Ponceau stain and rinsed with dH20 to visualize protein banding, then cut at 55kD into two pieces for separate antibody incubation. Membrane pieces were rinsed with dH20 to remove any residual Ponceau, then blocked for 1 hour in 1xTBS (Quality Biological 351-086-101) with 5% milk. Membrane pieces were incubated in antibody solution {1xTBS with 1% milk and 0.2% Tween 20 (Sigma P7949)} with primary antibodies against GAPDH and CRMP2 **(Supplementary Table 3)** overnight at 4°C. The next day membranes were washed four times with 1xTBST for 5 minutes with gentle shaking, followed by incubation with secondary antibodies **(Supplementary Table 3)** for 1 hour at room temperature with gentle shaking. Membranes were washed another 4 times with 1xTBST, then once with 1xTBS before imaging on a Licor Odyssey Clx with Image Studio 3 software. Images were quantified with Image J.

For the neuron blot, bands between 64 and 72kD were interpreted to represent CRMP2-B with or without possible post-translational modifications. Similarly, bands between 72 and 85kD were interpreted to represent CRMP2-A. This is because our CRMP2 antibody has previously detected CRMP2-B at 64kD and CRMP2-A at 72kD using the same ladder^14^ ; and because the lysis buffer we used is intended to preserve post-translational modifications including phosphorylation. When we did an additional Western Blot using the same reagents except with RIPA lysis buffer (which does not maintain phosphorylation), we observed the expected two bands at 64 and 72kD (data not shown). According to the RNA-seq data *DPYSL2*-*C* was not expressed, so we did not consider any protein bands to represent this isoform.

### Immunocytochemistry and dendrite morphology analysis of glutamatergic neurons

For dendrite morphology analysis, glutamatergic neurons were differentiated on a layer of rat astrocytes as described above. On DIV 21 immunocytochemistry was performed, with all washes and incubations at 4°C with no shaking using 300uL of cold reagents per chamber unless otherwise specified. The neuronal media was removed and chambers were rinsed with cold 1xPBS, then fixed with cold 4% paraformaldehyde (Alfa Aesar J61899) for 30 minutes. Samples were rinsed once then washed 3 times with cold 1xPBS; blocked in 1xPBS with 5% goat serum (Cell Signaling Tech. 5425S) and 0.3% Triton X-100 (Sigma 93443-100mL) for 1 hour; then incubated overnight with appropriate primary antibodies (**Supplementary Table 3**) in antibody solution consisting of 1xPBS with 1% BSA (Millipore Sigma A4161-1G) and 0.3% Triton X-100. The next day samples were rinsed once then washed three times with 1XPBS; incubated at 37C for 1 hour with appropriate secondary antibodies (see **Supplementary Table 3**) in antibody solution; then rinsed once and washed three times with 1XPBS. After removing the chambers from the slides, glass cover slips (Fisher1254M) were mounted with 25uL of Prolong Gold antifade reagent with DAPI (Invitrogen P36931) per chamber. After curing at 4°C for 24 hours slides were imaged on a Zeiss Axio Observer with Zen Blue software. Ten images at 4x magnification were captured for each of the 6 samples. The images were then blinded and the average length of the dendrites per cell was determined for each image using the MAP2 and NEUN channels in Image J (v 1.53) with the Neurphology J plugin. The images were then unblinded and the average dendrite length per cell was compared between the frameshift (array 1) and control (array 2) cells with a student’s t test. We did not use antibodies against CRMP2 because *DPYSL2* is expressed in astrocytes.

### Transcriptome analysis

Six frameshift and six control neuron samples were submitted to Novogene for paired-end RNA-sequencing. The twelve libraries were sequenced in one batch and all passed Novogene’s post-sequencing quality control. We received on average 25.3 million reads per sample with a maximum of 25.8 and a minimum of 24.3 million. The 150bp paired-end reads were aligned to human reference genome GRCH38 by Hisat2 version 2.1.0^47^. Samtools 1.9^48^ produced corresponding BAM files and Stringtie 2.0.3^49^ was used to assemble and estimate the abundance of transcripts based on GRCh38 human gene annotations^50^. Bioconductor package tximport computed raw counts by reversing the coverage formula used by Stringtie with the input of read length^51^. The output then was imported to another Bioconductor package DESeq2 for differential gene expression analysis^52^. Benjamini-Hochberg adjustment was used to calculate padj.

#### Gene Set Enrichment Analysis

expression files (.gct) were created using the counts function of DESeq2. Permutation type was set to gene set due to having less than 7 samples per group (frameshift vs controls). For custom annotation sets, topics were selected based on known *DPYSL2* functions from literature. Keywords corresponding to each topic (indicated in **Table 2** in quotation marks) were searched in the MSigDB XML Browser in version 7.4 and exported as gene set matrix files for input into GSEA. For pre-built annotation sets the default min and max gene set values were used (15 and 500 respectively). For custom annotation sets, the min and max gene set values were adjusted to accommodate all gene sets.

**Table 2.** Functional Classification of DESeq2 significant genes (padj < 0.05)

| Category | Number of Genes | Genes |
| --- | --- | --- |
| Neurotransmission / Neurodevelopment | 14 | *GRIA2, PIP5K1B, PHOX2A, EMX2, MEG3, BHCE, CHN1, PRDM8, TMPRSS15, OR6M1, C093525.2 (TBC1D24 and ATP6V0C), AC008758.1 (ZNF709 and ZNF564), Z84488.2 (CALHM6), AC025165 (AVIL)* |
| Microtubule/ Cytoskeletal dynamics / Cell Migration | 10 | *NEAT1, RBFOX1, PIP5K1B, SPAG6, FLG, SSTR2, PLXNA2, DYNC1H1, TLE4, AC092143.1 (TUBB3)* |
| Calcium Signaling/ Cell Adhesion | 8 | *ADGRV1, FAT1, SSTR2, PIP5K1B, CLEC4C, RBFOX1, Z84488.2 (CALHM6), AC244517.4 (PCDHB16)* |
| Immune response / Inflammation | 9 | *NLRP2, HS3ST4, KLRC4, CLEC4C, KLRC4-KLRK1, TXNIP, AL137247.2 (N4BP2L1), Z84488.2 (CALHM6), PPAN-P2RY11* |
| Protein Transport/ Vesicle trafficking | 4 | *GOLGA8A, PIP5K1B, ABCB11, AC093525.2 (TBC1D24),* |

#### GWAS Analysis for enrichment in DESeq2 significant genes

We downloaded all ensembl transcripts and their coordinates (hg19) from the University of California San Diego genome web site. These were merged into genes with their start and end position corresponding to the furthest extending transcripts. We downloaded the results of the largest schizophrenia GWAS to date ^53^from the Psychiatric genomics consortium at https://pgc.unc.edu/for-researchers/download-results/ (file name PGC3_SCZ_wave3_public.v2.tsv.gz) and filtered them to retain all SNPs with a p-value less than 1×10-7. This threshold which is slightly higher than genome wide significance (5×10-8) was used to maximize power while maintaining a strong enrichment for true signals. Those were then assembled into linkage disequilibrium groups that were at least 500kb apart from each other. This resulted in 266 such peaks of which only 31 were greater than 5×10-8. We then identified all ensembl genes that were within or less than 500kb from either direction of the linkage disequilibrium groups and considered all these genes as likely to drive the association. Enrichment was assessed by chi-squared test.

#### STRING analysis of DESeq2 significant genes

the ensembl IDs of *DPYSL2* and the 138 DESeq2 genes with padj < 0.2 were uploaded as a list to STRING ^54^ (https://string-db.org/). Of these 139 genes, 97 had protein products and were used for analysis. Enrichment of protein-protein interactions was calculated by STRING ^55^ using the whole genome as the statistical background. To further investigate functional enrichments, we added one 1^st^ shell interactor in the basic setting menu, which ended up being *PIK3R*. We also uploaded the 101 upregulated genes (67 protein products) and 37 downregulated genes (29 protein products) as separate lists.

#### Literature search of differentially expressed genes

we searched Pubmed for connections between our genes of interest and *DPYSL2*/CRMP2/mental illness by using the search command “Gene_name AND search_term”. The search term was either “*DPYSL2*”, “CRMP2”, or “” Mental Disorders”^53^ ”.

#### Enrichr Analysis

we uploaded the 75 DESeq2 genes (Entrez ID) at padj <0.05 to the Enrichr database (https://maayanlab.cloud/Enrichr/) as two lists: one of the 17 downregulated genes, and one of the 58 upregulated genes. We compared these lists to the “Old CMAP up”, “Old CMAP down”, and “PhenGenI Association 2021” libraries.

## RESULTS

### CRISPR editing and characterization of iPSCs

In order to exclusively knock out *DPYSL2*-*B*, we targeted its unique first exon **(Fig. 1a)**. Using CRISPR/Cas9 we generated six iPSC clones with a 1bp insertion (p.Iso12AsnC->CT) in exon 1 predicted to cause a frameshift and a premature termination codon in exon 2 **(Fig. 1b)**. qRT-PCR analysis confirmed this prediction by demonstrating a 3-fold reduction of *DPYSL2*-*B* expression in the frameshift iPSC clones relative to controls, consistent with the occurrence of nonsense-mediated decay **(Fig. 1c)**. As expected, CRMP2-B protein was also drastically reduced in frameshift clones **(Fig. 1d,e)**. To our knowledge, *DPYSL2*-*A* and *DPYSL2*-*C* are not expressed in iPSCs ^56^ and we saw no evidence of CRMP2-A at the expected sizes of 72kD ^14^ in our Western blot. Edited iPSC clones were negative for off-target CRISPR editing screened by targeted Sanger sequencing at the single candidate site that passed our prioritization pipeline. The edited and unedited iPSC lines were also negative for chromosomal abnormalities assayed via SNP array.

**Figure 1.**
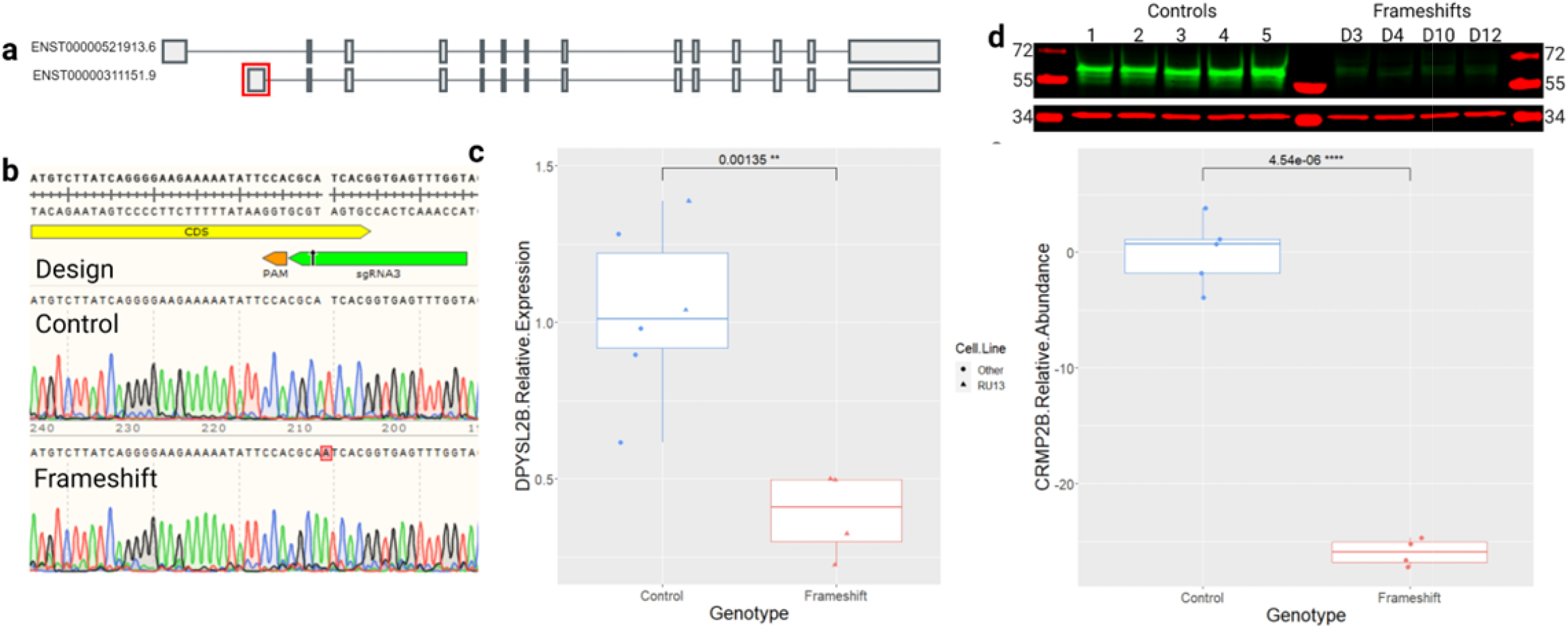
CRISPR Design and Editing in iPSCs. **(a)** DPYSL2 isoforms A and B (top to bottom). The first exon of DPYSL2B (boxed in red) was targeted to avoid impacting other transcripts. Image adapted from the GTEx portal. **(b)** Top panel shows the sgRNA (green) and PAM (orange) used to create the frameshift mutation in the coding sequence (CDS, yellow) in exon 1. Bottom two panels show Sanger sequencing traces of DNA from control and frameshift iPSCs. **(c)** qRT-PCR results showing significantly reduced expression of DPYSL2B in frameshift iPSC clones (n=4) compared to control clones (n=6 including 5 cell lines) (p=0.001, t test). Data were normalized to the geometric mean of GAPDH and B-actin loading controls, then to the average of the control clones. **(d)** Western blot comparing CRMP2B (∼64kD, green) abundance in control (n=5) and frameshift (n=4) iPSC clones with GAPDH (∼37kD, red) as a loading control. **(e)** Quantification of **(d)** showing significantly reduced abundance of CRMP2B in frameshift iPSCs compared to controls (p=4.5 × 10^−6^, t test). Data were normalized to GAPDH and then to the average of the control clones.

### Generation and characterization of excitatory glutamatergic neurons

Next, we used an *Ngn2*-induced differentiation protocol ^57-59^ with six frameshift and six unedited control iPSC clones to generate excitatory glutamatergic neurons **(Fig. 2a)**, a cell type repeatedly implicated in schizophrenia etiology ^18; 60; 61^. RNA-sequencing of these twelve clones showed that the neurons expressed appropriate neural and glutamatergic markers, and that the frameshift mutation did not significantly alter expression of these markers, i.e., the identity of the cells **(Supplementary Table 4)**.

**Figure 2.**
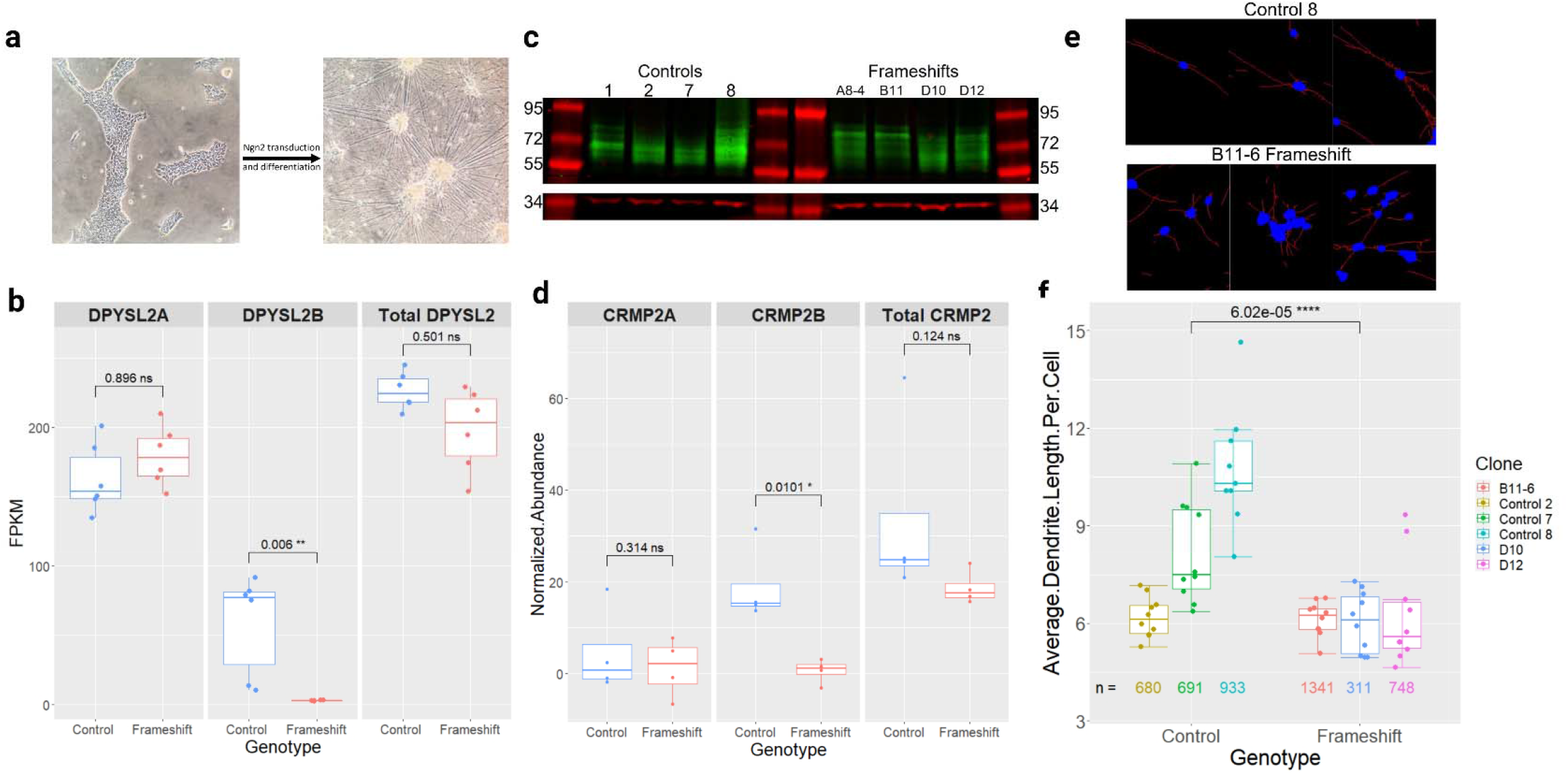
Generation and Characterization of DPYSL2B frameshift glutamatergic neurons. **(a)**Representative images of iPSCs before and after Ngn2 neuronal differentiation. **(b)** Quantification of RNA-seq data showing significantly reduced expression of DPYSL2B in frameshift (n=6) neuron clones compared to controls (n=6) (padj = 0.006, Wald test with Benjamini and Hochberg adjustment). DPYSL2A expression was not significantly changed, and all other CRMP2 isoforms were not expressed. **(c)** Western blot comparing CRMP2B (64-72kD, green) and CRMP2A (72-80kD, green) abundance in control (n=5) and frameshift (n=4) neuron clones with GAPDH (∼37kD, red) as a loading control. See the methods section for more detail on isoform distinction. **(d)** Quantification of **(c)**. CRMP2B was significantly reduced in the frameshift clones (p= 0.01, t test), but CRMP2A and the total amount of CRMP2 were not changed. Data were normalized to GAPDH. **(e)** Representative images of control and frameshift neurons after Neurphology J analysis with soma in blue and dendrites in red. Each image is cropped from the full-size pictures in Supplementary Figure 1. **(f)** Quantification of the average dendrite length per cell in frameshift (n=3) and control (n=3) neuron clones. Each dot represents data from one image, with 9-10 images analyzed per clone. The numbers under each boxplot represent the total number of cells that were analyzed for that clone across all the images. The average length of dendrites per cell was significantly reduced in the frameshift clones (p= 6.02 × 10^-5, t test).

#### Expression analysis of DPYSL/CRMP family and protein analysis of CRMP2

Our RNA-seq data showed that as expected, *DPYSL2*-*B* expression was significantly reduced in the frameshift neurons relative to control neurons. *DPYSL2*-*A* expression was higher than *DPYSL2*-*B* across both groups and was slightly elevated in the frameshift neurons, but this change was not significant. At the gene level, *DPYSL2* was significantly differentially expressed nominally but not after adjustment; we attribute this to the fact that most of the reads came from *DPYSL2*-*A* **(Fig. 2b)**. The remaining *DPYSL2* isoforms listed in ensembl were either not expressed (all samples had FPKM < 1; ENST00000523027, ENST00000493789, ENST00000521983, ENST00000474808) or not queried by the RNA-seq (ENST00000523690, ENST00000523093). The expression of *DPYSL*/*CRMP* –*1*, -*3*, -*4*, and –*5* were not significantly changed at the gene level. Analysis by isoform of the expressed transcripts found differential expression of one CRMP1 isoform (CRMP1-204, padj = 0.007) and two DPYSL3 isoforms (DPYSL3-201, padj = 0.042 and DPYSL3-202, pnom = 0.015, padj = 0.256). However, it should be noted that the CRMP1 isoform was very lowly expressed, and the fold change of the DPYSL3 isoforms was small (0.51 and 1.23, respectively) **(Supplementary Fig. 1)**.

At the protein level, as anticipated CRMP2-B was significantly downregulated in the frameshift neurons compared to controls. Interestingly, this reduction was not as profound as it was in the iPSCs (∼26x lower in iPSCs compared to ∼18x lower in neurons). CRMP2-A and the total abundance of CRMP2 did not change meaningfully between the two genotypes **(Fig. 2c)**. We noted that the abundance of CRMP2-A was lower than CRMP2-B in all the neuron samples, even though the expression of the *DPYSL2*-*A* transcript was higher than the *DPYSL2*-*B* transcript.

We next asked if knockout of endogenous *DPYSL2*-B is sufficient to cause aberrations in dendrite morphology. To investigate this, we differentiated three frameshift and three control clones on rat astrocytes to reduce soma clustering and performed immunocytochemistry with antibodies against the neuronal nucleus marker NEUN and the dendrite marker MAP2 **(Fig. 2e, Supplementary Fig. 1)**. We observed wide variation in the average length of dendrites in the control neurons; but quantitative analysis demonstrated a significant reduction of up to 58% of the average length of dendrites in the frameshift neurons compared to the controls **(Fig. 2f)**.

#### Transcriptome analysis of *DPYSL2*-B knockout glutamatergic neurons

To investigate how knockout of endogenous *DPYSL2*-*B* impacts the neuronal transcriptome, we analyzed our RNA-seq data with two pipelines: Gene set enrichment analysis (GSEA) ^62^ to identify expression changes of gene sets representing biological pathways; and DESeq2 ^52^ to identify differential expression of individual genes.

Agnostic GSEA using the broad “Hallmarks” and “GO Cellular Component” annotation sets revealed strongly significant (padj ≤ 0.05) downregulation of gene sets involved in mTORC1 signaling, protein secretion, myc targeting, and particularly the immunoglobulin complex; and upregulation of gene sets involved in the epithelial mesenchymal transition. Hypothesis-based GSEA with custom gene sets identified significant downregulation of gene sets related to calcium biology (padj ≤ 0.05). When we used slightly more permissive parameters of padj < 0.1, we also saw significant downregulation of gene sets related to cholesterol biosynthesis **(Table 1)**.

DESeq2 analysis with differentiation batch **(Supplementary Table 2)** as a covariate identified 75 genes with padj < 0.05 and 138 genes with padj < 0.2, most of which were upregulated **(Fig. 3a, Supplementary Tables 5 and 6)**. Of the 75 genes with padj <0.05, only four have been mentioned in Pubmed literature with *DPYSL2*/CRMP2 before: *CHN1*, encoding the signaling protein α2-CHN which is thought to act upstream of CRMP2 in the regulation of ocular motor axon guidance ^63^; IRS1, encoding a growth factor receptor which is part of the mTOR pathway that regulates CRMP2 ^64^; PLXNA2, encoding a Semaphorin 3A receptor which binds with CRMP2 and promotes its phosphorylation ^65^; and SSTR2, encoding a somatostatin receptor shown to mediate CRMP2 phosphorylation by regulating Ca2+ homeostasis ^66^. 24 of the 75 genes, including half of the top 20 most significant hits, were unnamed transcripts or novel lncRNAs with little to no characterization at the time of publication **(Supplementary Table 6)**. Analysis using the protein-protein interaction database STRING ^54^ with the 138 genes at padj < 0.2 at medium confidence and no additional interactors demonstrated a significant enrichment of interactions between their protein products **(Fig. 3c, Supplementary Table 6)**. To investigate any functional enrichments of these protein products we added a single 1^st^ shell interactor (STRING automatically selected PIK3R2). We observed enrichments in the KEGG diabetes mellitus type II pathway and the Monarch ^67; 68^ Intelligence and Cognition phenotypes. Separate analysis of the upregulated (101) and downregulated (37) genes also revealed a significant enrichment of upregulated protein products in the GO Cellular Component term Integral component of plasma membrane (GO:0005887, FDR = 0.043)**(Supplementary Table 6)**.

**Figure 3.**
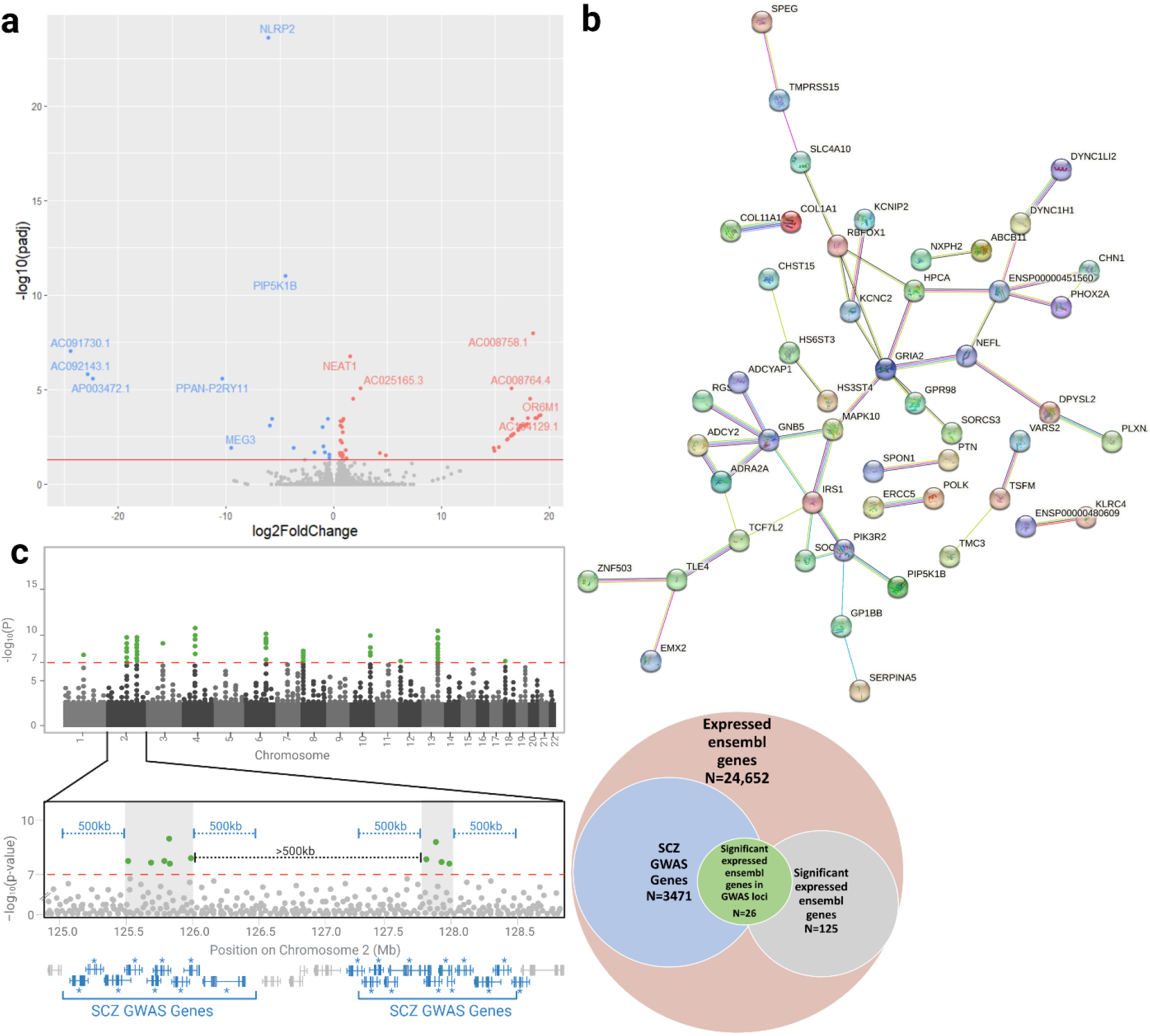
Transcriptomic analysis of frameshift neurons. **(a)** Volcano plot depicting differential gene expression of the frameshift and control neurons based on DESeq2 analysis of RNA-seq data. The red line indicates genome-wide significance of padj <0.05. Genes significantly upregulated in the frameshift neurons are in red and downregulated genes are in blue. **(b)** Schematic of theoretical GWAS data demonstrating the workflow used to identify genes in schizophrenia risk loci (“SCZ GWAS genes”) from the largest schizophrenia GWAS to date [35396580]. SNPs reaching significance of 1x 10-7 (green) were clustered into linkage disequilibrium blocks (gray shaded boxes) at least 500kb apart. Any genes within 500kb of these blocks (blue) were considered SCZ GWAS genes. Of the 24,652 genes expressed in the Ngn2 neurons, 3,471 were GWAS genes. Of the 125 genes that were present in the UCSC gene list and were significant with padj < 0.2 via DESeq2, 26 were GWAS genes. This represents a significant enrichment of our differentially expressed genes in schizophrenia risk loci (p=0.03, chi squared test). **(c)** STRING network demonstrating significant enrichment of protein-protein interactions amongst *DPYSL2* and the 132 differentially expressed genes with padj < 0.2. Of the 132 genes, 96 had protein products and 37 of these proteins interacted, representing a 1.77x enrichment (p=0.012, see methods for details on statistical analysis).

Review of the 75 differentially expressed genes highlighted *DPYSL2*’s role in the etiology of mental illness. The functional categories into which many of these genes fall are highly relevant to neuronal function **(Table 2)**. A Pubmed search of the 75 genes with the MeSH term “Mental Disorders” found that 42/75 are genes or read-through transcripts of genes already linked to neurological and/or psychiatric disease **(Supplementary Table 6)**. A parallel analysis also demonstrated that the 138 genes at padj <0.2 were enriched for localization in loci implicated in schizophrenia risk by GWAS. Specifically, we compared the 125 genes that had padj < 0.2 and were in UCSC’s hg19 gene list, and the 3,471 genes that are within 500kb of schizophrenia GWAS loci and were expressed in our neurons (24,652 genes). We observed that 26 genes overlapped whereas only 17.5 were expected, representing ∼50% more than expected (chi-square test 1.48x enrichment, p = 0.03; **Fig. 3c, Supplementary Table 7)**. Finally, we used the Enrichr database to compare the transcriptome profile of the 75 genes at padj < 0.05 to libraries from CMAP (https://www.broadinstitute.org/connectivity-map-cmap), as we had done in our work on HEK293 cells ^14^, and the Phenotype-Genotype Integrator (PhenGenI - www.ncbi.nlm.nih.gov/gap/phegeni). CMAP analysis showed that the 58 upregulated genes were significantly enriched for overlap with genes disrupted by the mTOR inhibitors sirolimus-1001 and LY-294002-6956 (p=0.003), the atypical antipsychotic sulpiride-1467 (p=0.034), and the calcium channel blockers tetrandrine-7178 (p=0.034). Comparison of the 17 downregulated genes with the PhenGenI database revealed overlaps with the atypical antipsychotic olanzapine (p=0.01) and the mood stabilizer lithium (p=0.054). These results show that knockout of *DPYSL2*-*B* disrupts genes and pathways involved in psychiatric disease.

A review of the most significantly differentially genes of the 75 further cements this. The most significantly differentially expressed gene in our dataset was the downregulated *NLRP*2, which encodes a protein key to the structure and function of astrocytic inflammasomes ^69^. *NLRP2* upregulation has been observed in bipolar disorder ^70; 71^ and familial early-onset Alzheimer’s, and suppression of *NLRP2* is the suspected mechanism of the candidate Alzheimer’s therapeutic GLP-1s ^72^. However, like in our dataset *NLRP2* has been shown to be downregulated in late-onset Alzheimer’s ^73^.

The second most downregulated transcript was AC092143.1 (ENSG00000198211), a novel protein-coding read-through transcript of *MC1R* and *TUBB3. TUBB3* (also known as TUJ1) encodes the neuron-specific class 3 beta tubulin, which heterodimerizes with other tubulins to form microtubules. As a main component of the neuronal cytoskeleton, TUBB3/TUJ1 is required for proper microtubule dynamics, cell migration, and axon formation. Deleterious mutations in *TUBB3* that interfere with heterodimerization ^74^ cause “tubulinopathies” which manifest with complex neurological and behavioral symptoms including cortical and cerebellar dysplasia, developmental delay, intellectual disability, cortical malformations, and reduced corpus callosum volume ^75-81^. Increased oxidative modifications of TUBB3 have been observed in Alzheimer’s brain tissue ^82^; and a recent study found reduced TUBB3 protein abundance in iPSC-derived organoids from schizophrenia patients ^83^. This same study also observed downregulation of CRMP1, CRMP2, and CRMP5 in these organoids. Although we did not observe downregulation of *TUBB3* itself in our dataset, we found it interesting that this read-through transcript was also downregulated.

Another downregulated gene of note was *MEG3*, a lncRNA tumor suppressor that is imprinted and maternally expressed. *MEG3* has been tied to a spectrum of neurological and psychiatric diseases, and recently has been investigated as a potential biomarker for these illnesses. Interestingly, the direction of *MEG3* expression varies between phenotypes and between studies. Some have shown that *MEG3* is upregulated in blood samples of children with autism ^84^, female schizophrenia patients ^85^, and in the brain of mouse models of Huntington’s ^86^ and Alzheimer’s ^87^. Other studies have shown that *MEG3* is downregulated in blood samples from male bipolar disorder patients ^88^, schizophrenia patients ^89^, and the hippocampus of rat ^90^ and mouse ^91^ models of Alzheimer’s and diabetes-related cognitive impairment ^92^.

Also among the top ten most downregulated transcripts were *PPAN-P2RY11*, a protein-coding read-through transcript downregulated in psoriasis ^93^ and whose parent gene *P2RY11* is downregulated in narcolepsy ^94-96^; and *PIP5K1B*, encoding a kinase that regulates actin dynamics ^97^, has been shown to be silenced in Friedreich’s ataxia ^98^, and SNPs in which have been associated with response to the atypical antipsychotic risperidone ^99^.

As for upregulated genes of note, *NEAT1* is a scaffolding lncRNA that aggregates with other RNAs in interchromatin spaces to form nuclear paraspeckles, which influence gene expression through their association with transcription, splicing, and epigenetic factors ^100^. *NEAT1* upregulation has been in observed in peripheral blood samples of patients with autism ^101^ and female patients with schizophrenia ^102^, as well in postmortem brain samples and mouse models of Huntington’s Disease ^103^. In Alzheimer’s disease, *NEAT1* is upregulated in the temporal cortex, hippocampus ^104^, serum and cerebrospinal fluid ^105^ and is thought to contribute to the pathology of Alzheimer’s by epigenetically influencing the activation of GSK3β, which phosphorylates Tau ^106^ and CRMP2. Also upregulated was *OR6M1* encodes an olfactory receptor and has very little literature; however, it was one of our SCZ GWAS hits. AC104129 (ENSG00000176349) is a novel lncRNA antisense to the gene *MAD1L1*, which encodes a mitotic spindle assembly checkpoint protein ^107^. GWAS have found associations between SNPs in *MAD1L1* and schizophrenia ^108-110^, bipolar disorder ^110-112^, and anxiety ^113^; and epigenomic changes in this gene are associated with post-traumatic stress disorder ^114; 115^ and schizophrenia ^116^. Finally, AC025165 (ENSG00000257921), a protein-coding read-through transcript of *EEF1AKMT*3 and *TSFM. TSFM* is a mitochondrial translation elongation factor and a homozygous variant in this gene has been reported to cause encephalocardiomyopathy and hearing loss ^117^.

## DISCUSSION

In the present study, we created and characterized the first iPSC and neuronal models of endogenous *DPYSL2*-*B* knockout. We used CRISPR/Cas9 in iPSCs to create a frameshift mutation in *DPYSL2*-*B*’s private first exon, which resulted in significant reduction of the mRNA and protein both in iPSCs and in *Ngn2*-induced excitatory glutamatergic neurons.

### DPYSL2-B knockout did not significantly alter expression of CRMP2-A

At the time of publication, *DPYSL2* has 8 transcripts listed in ensembl. RNA-seq data from a recent study by our lab and others ^56^ of nine iPSC lines showed that *DPYSL2*-*B* is the only isoform expressed in iPSCs, so there were no other transcripts to query for differential expression. As for neurons, here we found that only *DPYSL2*-*B* and *DPYSL2*-*A* were expressed in Ngn2-induced neurons, a pattern consistent with other Ngn2 datasets we’ve generated ^58^. We observed that knockout of *DPYSL2*-*B* did not significantly change expression of *DPYSL2*-A/CRMP2-A **(Fig. 2b-d)**. Others have found that these two isoforms have opposing effects on axon elongation ^36^ and suggested they may work in tandem. We did not observe any significant compensatory reaction at the transcript or protein level, and our neuron cultures were too dense to assay axon morphology. Regarding other functional overlaps, our gene set enrichment analysis revealed that knockout of *DPYSL2*-*B* resulted in upregulation of genes related to the epithelial-mesenchymal transition, an oncogenic process that enables metastasis. This process was found to be suppressed by CRMP2 ^118^, and later more specifically by CRMP2-A ^41^. Our GSEA result suggests the epithelial-mesenchymal transition may also be regulated by CRMP2-B, highlighting a potentially novel functional overlap between these two isoforms. Overall, our results suggest that if CRMP2-A does indeed have an opposing effect to CRMP2-B, it does not appear to be mediated by compensatory changes in *DPYSL2*-*A* RNA or protein expression.

### Knockout of DPYSL2-B disrupts dendritic morphology and pathways involved in cytoskeletal dynamics

We found a significant reduction of the average length of MAP2+ dendrites in our frameshift glutamatergic neurons **(Fig. 2e-f)**. In previous work when we reduced translation of the *DPYSL2*-*B* isoform by editing the TOP sequence in the 5’ UTR in HEK293 cells we observed a marked reduction in the length of their projections ^14^. Shortening and/or aberrant morphology of neurites has also been observed in *DPYSL2* knockout mouse models that disrupt all *DPYSL2* isoforms ^8; 24; 25; 119^, as well as in rat hippocampal neurons transfected with *DPYSL2*-*B* truncation mutant transgenes ^38^. Our findings in tandem with others confirm that the *DPYSL2*-*B* isoform is necessary for proper dendrite formation.

Additionally, our DESeq2 analysis identified differential expression of several genes that compose, regulate, and utilize the cytoskeleton **(Table 2)**. *AC09214*3 is a read-through transcript of the neuron-specific tubulin *TUBB3* (discussed above). *SPAG6* encodes a microtubule associating protein crucial to the function of the axeome central apparatus, a microtubule structure in flagella and cilia ^120^. *FLG* is an actin filament-associating protein recently shown to regulate BACE1, an APP-cleaving enzyme that contributes to AB plaque formation in AD ^121^. Several genes regulate the kinase activity of GSK3β upon CRMP2, including *NEAT1* (discussed above), *TLE4* (a transcriptional co-repressor regulated by Wnt ^2; 122^), *PLXNA2* (a Semaphorin 3A co-receptor, discussed above), *PIP5K1B* (a kinase that regulates the production of PI3K ^97; 98^) and *SSTR2* (co-receptor for somatostatin, discussed above ^123^). *RBFOX1* (a neuron-specific splicing factor) regulates many neuronal development genes including those related to the cytoskeleton ^124; 125^. *DYNC1H1* encodes a heavy-chain component of cytoplasmic dynein, a motor protein complex that performs intracellular transport along microtubules. CRMP2 is known to directly bind to the heavy chain region of dynein and modulate its activity ^126^. *GOLGA8A* encodes a member of the Golgin protein family and is a blood biomarker for ASD ^127^. Among other functions, Golgins act as tethers between the cytoskeleton and Golgi. Golgins can also bind to dynein motors to facilitate transport of Golgi vesicles during mitosis ^128^. *ABCB11* belongs to a family of ubiquitous ATP-powered transporters that traffic cargo across intra and extra-cellular membranes ^129; 130^. The differential expression of these molecular transport-associated genes was corroborated by GSEA, which found downregulation of genes related to protein secretion. Taken together, these results confirm that *DPYSL2*-*B* is necessary for the development of the cytoskeleton, disruption of which leads to aberrations in neuronal structure and function that are highly relevant to psychiatric disease.

### Involvement of DPYSL2-B in neuronal cholesterol biosynthesis

Our GSEA revealed that knockout of *DPYSL2*-*B* in glutamatergic neurons downregulated the expression of genes involved in cholesterol biosynthesis. STRING analysis of our upregulated DESeq2 genes also found an enrichment of proteins integral for plasma membrane structure. As a major component of myelin, cholesterol is imperative for proper insulation of axons for saltatory conduction of electrical impulses along axons. It is also an essential structural component of the plasma membrane; a substructure which is both particularly abundant in neurons due to the large surface area created by their dendrites, axons, and terminal branches; and particularly functionally critical for the proper transmission of neurotransmitters and electrical impulses from cell to cell.

Dysregulation of lipid homeostasis has been implicated in the etiology of numerous brain disorders including schizophrenia, Alzheimer’s, Parkinson’s, and Huntington’s disease ^131; 132^. Postmortem analysis of brain tissue from patients with schizophrenia has found reduced myelination ^133^, as well as decreased levels of phospholipids and polyunsaturated fatty acids ^131^. Additionally, multiple studies have reported that antipsychotics modulate lipid homeostasis and suggested that this contributes to their therapeutic effects ^131; 134^. This mechanism of action makes sense considering antipsychotics commonly have metabolic side effects including weight gain and increased insulin resistance ^135^. Of particular relevance to schizophrenia, our lab recently demonstrated that treatment of *Ngn2* neurons with clozapine results in upregulation of cholesterol synthesis genes ^58^. When we compare those upregulated genes to our current study, we see that 6/7 of our GSEA leading edge genes match, except our genes were downregulated (*HMGCS1, IDI1, HMGCR, FDFT1, FDPS, CYP51A1*). Similarly, when we compared the transcriptome profile of our 17 DESeq2 downregulated genes at padj <0 .05 to those of ConnectivityMap perturbagens, our #2 hit was olanzapine, an atypical antipsychotic. The fact that our gene expression patterns are complementary to those of antipsychotics confirms that knockdown of *CRMP2-B* captures schizophrenia disease signal, and further cements the link between schizophrenia, lipid homeostasis, and *DPYSL2*-*B*/CRMP2 function.

At the molecular level, the differential expression of cholesterol biosynthesis genes could indicate disruption of lipid structures that mediate Semaphorin 3A signaling. Within the plasma membrane cholesterol is enrichment within lipid rafts, which act as microdomains that facilitate the aggregation and interaction of extracellular signaling molecules and their receptors, thereby encouraging propagation of intracellular signaling cascades ^136^. Several studies have shown that components of the Semaphorin 3A signaling cascade interact with lipid rafts ^137^. This includes the Semaphorin 3A receptor plexin A2, the gene for which (*PLXNA2*) was significantly upregulated in our frameshift neurons. Two studies have shown that in the rodent brain CRMPs themselves can co-localize with lipid rafts ^137; 138^. Furthermore, studies have shown that some lipid rafts can be tethered to the cytoskeleton, which would provide an even more direct interface for cytoskeletal-associated proteins like CRMPs and their upstream regulatory effector molecules ^137^. Taken together, this data supports the notion that the disruption of cholesterol synthesis could impact how CRMP2-B is regulated by its upstream effectors.

It is also possible that CRMP2 itself could be regulating cholesterol synthesis. A recent study by Chang et al. found that CRMP2 can regulate lipid metabolism through multiple mechanisms during adipocyte differentiation in the mouse brain. Interestingly, they found opposite results to ours. Overexpression of CRMP2 via transgene led to downregulation of lipid metabolism; and conversely, knockdown by CRMP2 siRNA led to increases in lipid accumulation ^139^.

### Knockout of DPYSL2-B confirms link between mTOR and schizophrenia

The molecular target of rapamycin, or mTOR, is a serine/threonine kinase that ubiquitously regulates processes such as proliferation, survival, and autophagy ^140; 141^. The mTOR complex mTORC1 regulates mRNA translation by phosphorylating effector proteins which influence binding of translational initiation factors to 5’ TOP motifs in the 5’UTR of mRNAs ^140^. In neurons, mTOR influences polarization and axon elongation by regulating translation of microtubule-associating proteins like Tau and CRMP2 ^15; 142^. mTOR dysregulation has been implicated in numerous psychiatric disorders ^143^. The SCZ-associated 13 DNR we previously studied is within a TOP motif in the 5’UTR of *DPYSL2-B*. This variant resulted in reduced binding of *DPYSL2-B* mRNA to mTOR effector proteins ELALV4 (ribosomal binding protein that transports mRNA from the nucleus to the cytosol), eIF4E (a translation initiation factor) and 4EBP (a translational repressor that inhibits eIF4E). We had also showed that HEK293 cells with the 13DNR were more sensitive to treatment with Rapamycin, an mTOR inhibitor, than controls ^14^. Here we confirm that the *DPYSL2*-*B* isoform is involved in mTOR signaling in glutamatergic neurons. Our GSEA found that knockout of *DPYSL2*-*B* results in downregulation of genes involved in mTORC1 signaling. These results are consistent with a recent study that observed downregulation of protein expression and phosphorylation of Akt/mTOR pathway proteins in the dorsal lateral prefrontal cortex of schizophrenia patients ^144^. DESeq2 identified significant upregulation of *IRS1*, a growth factor receptor upstream of mTORC1 in the PIK3/Akt/mTOR pathway. Furthermore, the 57 genes upregulated in the frameshift neurons had a transcriptome profile complementary to mTOR inhibitors sirolimus and LY-294002-6956. These results further confirm the role of the *DPYSL2*-*B* isoform as a common link between mTOR and schizophrenia.

### DPYSL2-B knockout results in disruption of immune system genes

Disruption of inflammatory pathways has long been realized to contribute to the risk of psychiatric disease, especially schizophrenia ^145^. Our strongest GSEA result was a downregulation of genes involved in the immunoglobulin complex. Deseq2 also identified several immune-related differentially expressed genes, including inflammasome components *NLRP2* (discussed above) and *TXNIP*, a thioredoxin-binding protein that regulates oxidative stress ^146^. *TXNIP* dysregulation is associated with numerous psychiatric disorders including Alzheimer’s and schizophrenia ^147-150^. We observed strong upregulation of *KLRC4*, its readthrough transcript *KLRC4-KLRK1*, and *CLEC4C*, all of which encode lectin receptors thought to play a role in antigen processing in the innate immune system ^151^, and the former two of which are dysregulated in the autoimmune disorder Behcet’s ^152^. These findings support the notion that *DPYSL2*-*B* plays a role in early immune system function and further links this system with psychiatric disease.

## Limitations and future directions

Our study has a few limitations that could be addressed by future work. Although we found significant reduction of *DPYSL2*-*B* RNA and protein in our *Ngn2*-induced neurons, our Western blot suggests that CRMP2-B was not completely ablated. The cells were genotyped prior to differentiation, so we are confident that the neurons were not contaminated with unedited cells. The presence of residual CRMP2-B protein suggests that there is some sort of amelioration of the frameshift mutation that we cannot yet explain nevertheless, its expression is greatly reduced. Our first approach to knocking out *DPYSL2*-*B* was to delete the majority of the first exon, which we successfully accomplished in iPSCs. However, we observed a consistent loss of the deletion genotype despite repeatedly purifying the population with multiple rounds of single-cell cloning (data not shown), which suggests that the deletion cells may have a growth disadvantage. Additionally, according to the genome aggregation database (https://gnomad.broadinstitute.org/) human nonsense variants in *DPYSL2* are exceedingly rare, are never observed as homozygotes, and never occur in this isoform prior to the 8^th^ exon. Consequently, *DPYSL2* is predicted to have a probability of loss of function intolerance (pLI) score of 0.99 out of 1, supporting the notion that it is an essential gene. It should be noted that homozygous *DPYSL2* knockout mouse models have been successfully generated ^8; 24; 25; 119^ yet one should consider the different organism and that there may be more buffering systems *in vivo* that could compensate for disruption of a critical gene.

In summary, we generated the first human iPSC and neuronal models of endogenous *DPYSL2*-*B* knockout. Our findings confirm the crucial role of this specific transcript in the proper development of the cytoskeleton and numerous pathways influencing neurodevelopment and related to psychiatric diseases including schizophrenia.

## ACKNOWLEDGEMENTS

This work was supported by NIMH grants R01 MH113215 and RF1 MH122936 to DA.

