## Supplemental material for "*DPYSL2*/*CRMP2* isoform B knockout in human iPSC-derived glutamatergic neurons confirms its role in mTOR signaling and neurodevelopmental disorders"

**Supplementary Figure 1 | Full size images of stained neurons post Neurphology J processing**

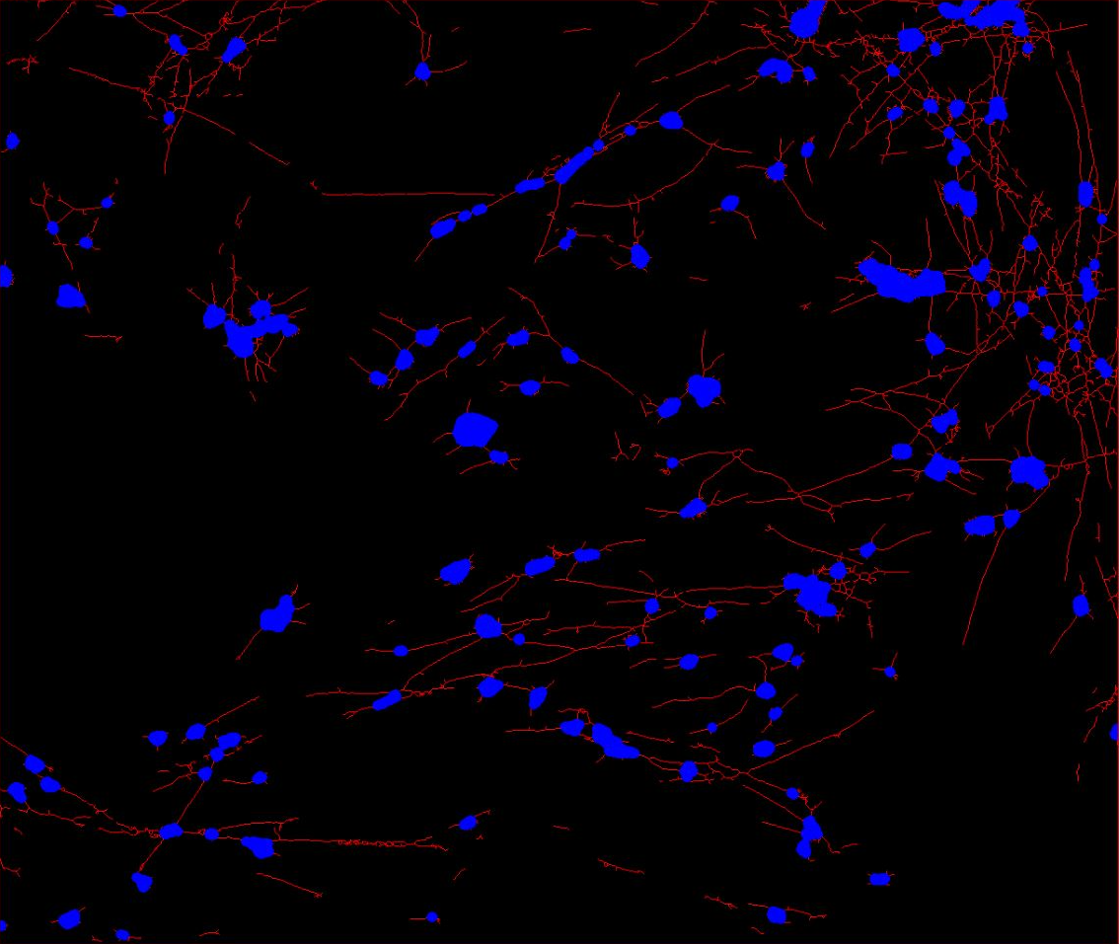
**B11-6 Frameshift**

**Control 8**

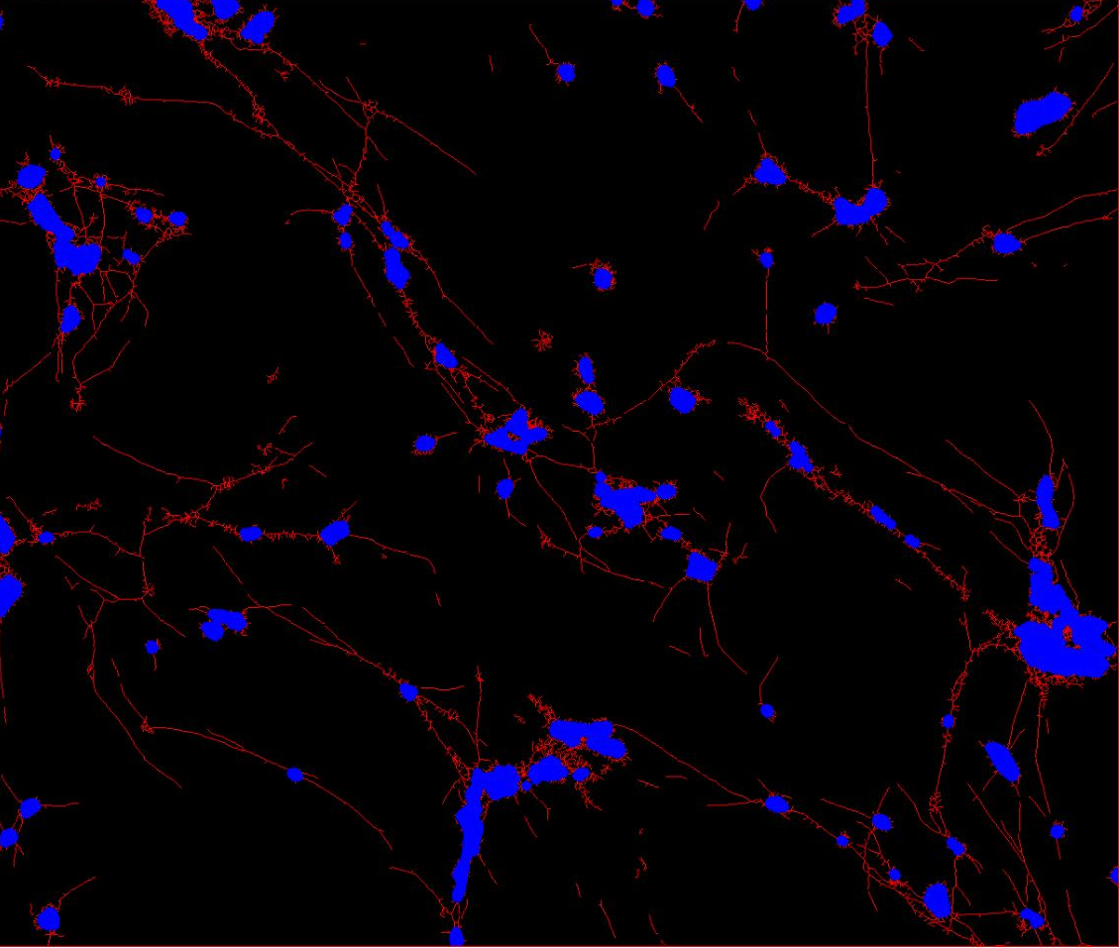

**Supplementary Information**

**perl script used to find overlaps between candidate CRIPSR off-target sites and open chromatin peaks/exons**

- **infile 1 should be a CRISPOR (http://crispor.tefor.net/) report of candidate off-target sites, the name and head of which should look like:**

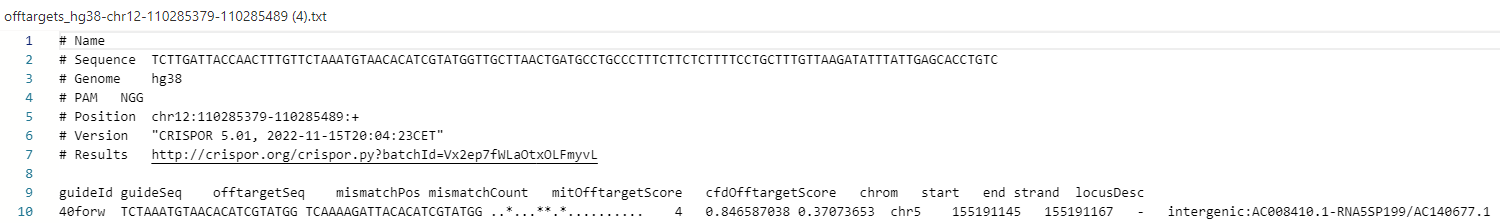

- **infile2 is an hg38 open chromatin bed file from encode (**[**www.encodeproject.org**](http://www.encodeproject.org)**, file name hg38_wgEncodeRegDnaseClustered.txt)**
- **set $distance to the number of bp on either side of the open chromatin peaks that you want to include**

my $infile1 = $ARGV[0] ;

my $infile2 = "hg38_wgEncodeRegDnaseClustered.txt"; #$ARGV[2] ;

my $distance = $ARGV[1] ;

my $outfile = "overlaps.$distance.$ARGV[0]";

my @chr = "";

my @start = "";

my @end = "";

open (INFILE2, "$infile2") or die "Unable to open $infile2 for reading\n";

my $count=0;

my $xx=1;

my $found=0;

$|=1;

print "\n\nreading chomatin marks input\n";

while(<INFILE2>){

chomp; my @line2 = split;

$chr[$count]=$line2[1];

$start[$count]=$line2[2]-$distance;

$end[$count]=$line2[3]+$distance;

$count++;

if($count/100000 > $xx){print "\r$xx"."00,000 lines read";$xx++;}

}

open (OUTFILE, ">$outfile") or die "Unable to open $outfile for writing\n";

print OUTFILE "fragments from $infile1 overlapping with fragments from $infile2 or within $distance bp:\n";

print "\n\ncomparing to off targets \n";

open (INFILE1, "$infile1") or die "Unable to open $infile1 for reading\n";

while(<INFILE1>){

chomp; my @line1 = split;

if($line1[0] eq "\#"){print OUTFILE "$_\n";next;}

if($line1[0] eq "guideId"){print OUTFILE "$_\n";next;}

if($_ =~ /exon/){print OUTFILE "$_\n";next;}

for($i=0; $i<$count; $i++){

if($line1[7] eq $chr[$i] and (($line1[8] < $end[$i] and $line1[8] > $start[$i])

or ($line1[9] < $end[$i] and $line1[9] > $start[$i]))){$found++;print OUTFILE "$_\n"; print "\rfound $found overlaps";}

}

}

print "\n\n";

close INFILE;

close OUTFILE;
